# Assortative mating may have facilitated rapid evolution of locally adaptive phenologies during a biological invasion

**DOI:** 10.64898/2026.10.08.757682

**Authors:** María José Gómez Quijano, Mia N. Akbar, Emily Crowson, Robert I. Colautti

## Abstract

Assortative mating for a trait under selection can increase the rate of adaptive evolution and limit introgression of detrimental alleles. We therefore hypothesized that assortative mating facilitated the rapid evolution of flowering phenology observed in the self-incompatible perennial *Lythrum salicaria* (purple loosestrife) as it invaded eastern North America. We characterized flowering schedules of 600 individuals representing 20 populations sampled along a latitudinal gradient of 10 degrees and grown in a common garden at mid-latitude. Consistent with our hypothesis, we found that flowering schedules were less synchronous in the mixed population of the common garden (*s* = 0.50) relative to the 20 locally adapted populations comprising it (*s* = 0.56 to 0.86), and flowering synchrony decreased towards higher latitudes of origin. Despite high synchrony, assortment potential (*ρ*) by growth and phenology varied by population (*ρ* = 0.03 to 0.61), but the predicted negative correlation between *s* and *ρ* was masked by among-individual fitness variation in the common garden. We show how fitness variation and asynchrony associated with locally adapted phenology can reduce the fitness impacts of maladaptive gene flow and accelerate the rate of adaptive evolution during invasion.

## 1. Introduction

The evolutionary trajectories of natural populations are shaped by the interplay of stochastic and adaptive processes (Dobzhansky, 1937; Mayr, 1963; Wright, 1931). Phenology is often under strong selection from both biotic interactions and abiotic environments because it encompasses a suite of key life-history traits that determine the timing, frequency and success of reproduction (Anderson et al., 2012; Austen et al., 2014; Weis et al., 2014). Even in the absence of selection, variation in the timing of reproduction can lead to assortative mating and lower effective population sizes, thereby influencing the strength of genetic drift (Crow & Felsenstein, 1968; Devaux & Lande, 2008). In plants, climate heterogeneity is often associated with phenological differences that alter reproductive synchrony, species interactions and mating opportunities (Fox, 2003; Ison & Weis, 2017; Parmesan & Yohe, 2003; Rivest et al., 2023). Together, these processes highlight phenology as an important mediator of eco-evolutionary dynamics via effects on genetic drift and natural selection.

If natural selection and genetic drift are the drivers of evolutionary change, assortative mating is the accelerator pedal. Nonrandom mating for any phenotypic trait can be quantified as the phenotypic correlation between mates (Fisher, 1919), with positive values indicating assortative mating and negative values indicating disassortative mating. The genetic effects of positive phenotypic correlations between mates, hereafter referred to as assortative mating resemble those of inbreeding (Crow & Felsenstein, 1968), but apply only to genetic loci associated with the assorting trait (Crow & Felsenstein, 1968; Wright, 1921). By reducing mating between phenotypically dissimilar individuals, homozygosity of loci associated with the assorting trait(s) increases and genotype frequencies change, causing departures from Hardy-Weinberg equilibrium without directly altering population allele frequencies (Breese, 1956; Crow & Felsenstein, 1968; Lande, 1977). Assortative mating can therefore increase the evolutionary response to selection by generating nonrandom associations among loci associated with the assorting trait that increase additive genetic variance (Wright, 1921), and by exposing recessive alleles previously masked in heterozygotes when homozygosity increases (Crow & Felsenstein, 1968).

Outcrossing plants are well known to assort by flowering time (Breese, 1956; Fox, 2003; Weis et al., 2005), because mating opportunities among individuals depend on the extent to which their flowering schedules overlap (Austen et al., 2014; Peters & Weis, 2019; Shelton et al., 2024). Phenological traits that influence the timing of reproduction can reduce the probability of allele exchange between individuals, thereby increasing population structuring and promoting differentiation (Devaux & Lande, 2008; Ison & Weis, 2017; Perini et al., 2020; Peters & Weis, 2019; Vines & Schluter, 2006; Wadgymar, Cumming, et al., 2015). In outcrossing plant populations, variation in flowering onset and duration reduces reproductive schedule overlap among individuals, increasing reproductive asynchrony and partitioning the population into smaller, temporally structured mating pools (May et al., 2023; Wadgymar et al., 2015; Weis & Kossler, 2004). If individuals also vary strongly in fitness, this temporal structuring can be further amplified (Shelton et al., 2024; Wadgymar, Austen, et al., 2015), as a small number of high-fitness individuals will disproportionately contribute to reproduction. Consequently, even in large census-size populations, reproductive asynchrony and temporal structure arising from assortative mating and fitness differences can reduce effective population size, increasing the strength of genetic drift, and promoting genetic differentiation (Bolnick & Kirkpatrick, 2012; May et al., 2023; Vines & Schluter, 2006).

Reproductive synchrony in plants is often approximated by similarity in flowering onset (i.e., Augspurger, 1983; Forrest et al., 2010; Rodríguez-Pérez & Traveset, 2016). However, measuring entire flowering schedules provides opportunities to characterize all potential mating opportunities within a population, yielding mathematically concise estimates of mating symmetry and assortment (Lyons & Mully, 1992; Weis et al., 2014). However, the extent of phenological assortment and flowering synchrony among natural plant populations remains largely unexplored. The few studies that have characterized individual flowering schedules reveal substantial variation within and among populations, representing potential for assortment by phenology (i.e. Akbar et al., 2025; Austen et al., 2014; Ison & Weis, 2017; Peters & Weis, 2019). Importantly, this variation is shaped in flowering phenology and reproductive success is shaped by genetic effects, environmental conditions and genotype-by-environment interactions, meaning that mating opportunities are not fixed properties of individuals but instead arise from how their phenotypes interact with local environmental conditions. Therefore, variation in flowering schedules may influence the evolutionary consequences of populations by determining which genotypes successfully reproduce, potentially limiting maladaptive gene flow while facilitating the spread of locally adapted beneficial alleles.

Common garden experiments provide a powerful framework for assessing these processes by measuring detailed flowering schedules and mating opportunities among individuals from multiple populations in shared environment (Akbar et al., 2025; Austen et al., 2014; Wadgymar, Austen, et al., 2015). By assembling individuals across geographic gradients into a common environment, common gardens can approximate the structure expected during early-invasion stages characterized with multiple introductions and substantial genetic and phenotypic variation (Colautti & Lau, 2016; de Villemereuil et al., 2016; Huxman et al., 2022). In this context, common gardens offer a unique opportunity to quantify the potential for phenological assortment and assess how assortative mating may have facilitated adaptive trait differentiation during early invasion and continues to shape differentiation in contemporary populations.

Here, we quantify flowering schedule synchrony and the potential for assortative mating potential in a common garden experiment featuring an artificially mixed population composed of 20 natural populations of *Lythrum salicaria*. We then evaluate how variation in flowering onset and total flower production have influenced flowering synchrony and assortative mating potential. By assuming that contemporary populations are locally adapted relative to the artificially mixed population, we test the hypothesis that assortment facilitated the rapid evolution of locally adapted flowering time clines in North American populations of *L. salicaria* (i.e., Akbar et al., 2025; Colautti & Barrett, 2013; Montague et al., 2008; Wu & Colautti, 2022). Here, we analytically decouple mating schedule shape from flowering time and fitness variation and apply methods developed by Weis and Kossler (2004), Weis (2005) and Weis et al. (2014), to estimate reproductive synchrony (*s*) and the phenotypic correlations among mates (*ρ*). We use these metrics to address the following questions: 1) Do mixed and locally adapted populations differ in flowering synchrony and phenotypic correlation between mates? 2) Does assortative mating potential vary across latitudinal gradients? And 3) how does variance in flowering onset and individual fitness influence flower synchrony and the potential for phenological assortative mating? Given the mixed nature of the common garden, we predict that flowering schedules in the artificially mixed population will be less synchronized, resulting in a higher potential for assortative mating, relative to the individual populations comprising it. Second, we predict an increase in synchrony and a decrease in assortment in populations sampled from higher latitudes where populations experience stronger stabilizing selection on flowering phenology. Third, we predict that local adaptation to the mid-latitude common garden site will increase mating opportunities in plants with intermediate flowering times, increasing their mating opportunities relative to early- and late-flowering individuals.

## 2. Materials and Methods

### 2.1 Study System

*Lythrum salicaria* is an insect pollinated, outcrossing and tristylous wetland perennial herb that is native to Eurasia (Stuckey, 1980). Herbarium records indicate establishment in eastern North America by the late 18th century (Thompson et al., 1987). While *L. salicaria* can produce multiple flowering stems, most of its reproduction is localized to a larger primary stem that bears dozens to hundreds of self-incompatible and tristylous, small purple to pink flowers (Figure 1A) (Cheung & Sattler, 1967; Stuckey, 1980). Rapid population growth is facilitated by high seed production (Thompson et al., 1987), with some individuals capable of producing up to 2.7 million seeds in a single growing season (McCaughey & Stephenson, 2000).

**Figure 1.**
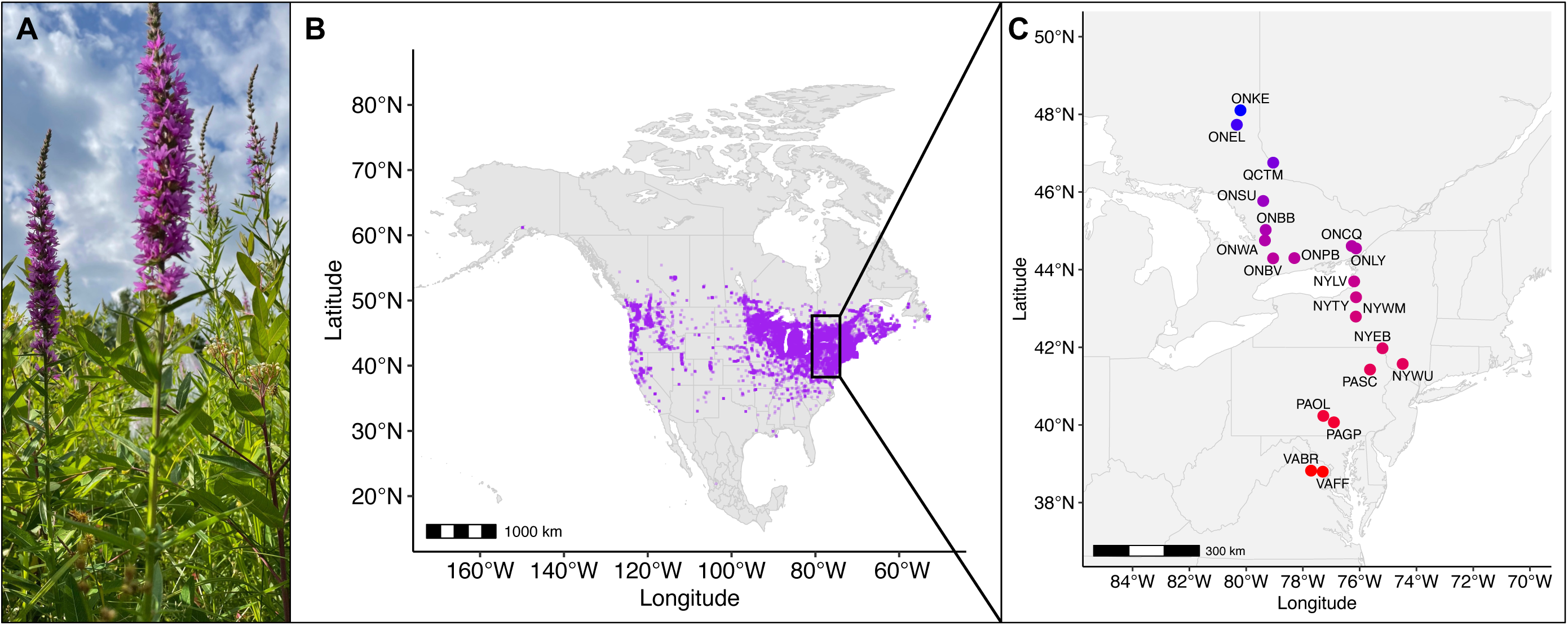
Overview of the study system: *Lythrum salicaria* in North America. A) Photograph showing primary stem and inflorescence of a *L. salicaria* individual grown in a common garden experiment at the Queen’s University Biological Station (QUBS). B) Map of occurrences (purple) of *L. salicaria* in North America from the Global Biodiversity Information Facility (GBIF). C) Map of eastern North America showing locations of 20 sampled *L. salicaria* populations spanning ten degrees of latitude. Study populations are identified by a four-letter code with the first two letters corresponding to the standard abbreviation of the U.S. state or Canadian province of origin. Populations are color-coded based on the latitude of origin with warmer tones representing lower latitudes and cooler tones representing higher latitudes.

The geographic range of *L. salicaria* remained localized to eastern North America with few records in Canada until the 1930s, when it experienced a rapid expansion across the continent between 30 ° to 50 ° N (Thompson et al., 1987), where it is now well established (Figure 1B) (Munger, 2002). Clines observed in multiple common garden experiments and an herbarium study support the hypothesis that earlier flowering at smaller vegetative sizes evolved at northern latitudes as an adaptive response to shorter growing seasons, with later flowering at a larger size in southern latitudes as a result of relaxed selection in longer growing seasons (Akbar et al., 2025; Colautti et al., 2010; Colautti & Barrett, 2013). A ‘virtual common garden’ analysis of herbarium records supports latitude-specific selection for flowering time across the North America range, and also revealed rapid cline evolution for the first 100 years of invasion, followed by a period of relative stasis (Wu & Colautti, 2022). To our knowledge, no studies have yet tested whether assortative mating could have contributed to the rapid evolution of flowering time clines in this study system.

### 2.2 Common Garden Experiment

We estimate potential for assortative mating based on flowering schedules measured in a common garden experiment established in 2017 at the Queen’s University Biological Station (QUBS) in Elgin, ON (44° 32’ 14” N 76° 22’ 25” W). The common garden was established using seeds collected in the fall of 2016 from 20 natural populations of *L. salicaria* spanning a 10-degree latitudinal gradient in eastern North America (Figure 1C), resampling along a gradient sampled almost a decade earlier (Montague et al., 2008). The common garden was situated near mid-latitude of this gradient, with 10 seed families from each of 20 populations (200 total) replicated across six randomized blocks. Three of six experimental blocks were exposed to herbivory from locally colonizing *Galerucella* beetles introduced for biological control and were not used in this experiment. The three remaining blocks were treated with deltamethrin in May of each year, prior to flowering, to limit herbivore damage. Thus, our experiment included 600 plants in total, with each of three experimental blocks containing the same 200 seed families representing 20 populations sampled along a 10-degree latitudinal gradient.

From June to September 2023, we recorded weekly measurements of the primary stem length, divided into sections containing only vegetative growth (i.e., no reproductive structures), buds, flowers, and fruits. We additionally monitored each plant daily to identify the date of first flower. Beginning on the first date of recorded flowering, we counted the total number of viable flowers for the entire plant every five days until flowering was complete. Flowers of *Lythrum salicaria* typically remain viable for less than 24 h after anthesis, with contraction of the calyx within 48 h as pollinated flowers begin to form fruits, or senescence of unpollinated flowers within five days (Waites & Ågren, 2006). Therefore, a census schedule of viable flowers every 5 days would avoid problems associated with recounting the same flowers on more than one census day.

### 2.3 Data Analysis

Of the initial 600 individuals, we removed 145 that did not produce at least 10 flowers over 10 days, and/or had damage to the primary stem. Our final dataset was comprised of 455 individuals across 20 populations, with 11 to 31 individuals per population. Our published dataset includes the date of first flower, the number of open flowers on each census day (5-day intervals), along with associated information on population location and size at flowering. It is important to note, that for this study, we assumed equal opportunity of mating among all individuals regardless of floral style morph. In *L. salicaria,* tristyly restricts mating to complementary style morphs and prevents self-incompatibility. However, due to sample size limitations and the complexity of the mating matrices, style was not incorporated into our analyses.

Because growth and phenology are highly correlated in this species, we used a scaled principal components analysis (PCA) to capture the correlation between day of first flower and length of the primary vegetative stem at flowering. We retained the first principal component (PC1) as a measure of phenotype, with positive values representing smaller plants that flower earlier and negative values representing larger plants that flower later. This is comparable to PC1 reported in several previous studies of *L. salicaria* examining the trade-off between flowering time and size at flowering in the context of natural selection and local adaptation (Akbar et al., 2025; Colautti et al., 2010; Colautti & Barrett, 2010, 2011; Wu & Colautti, 2022).

We used methods described in Weis (2005) and Weis and Kossler (2004). Briefly, we constructed a pairwise mating probability matrix (Φ), with each value representing the probability that a randomly chosen offspring resulted from a mating between a given pair of individuals. The sum across all values of Φ is equal to 1, with the diagonal representing self-pollination probabilities. The square matrix Φ is calculated as the product of two matrices: an M matrix of individuals (rows) and the proportion of receptive flowers open on each census day (columns) and a P matrix of individuals (columns) and the proportion of pollen-producing flowers on each census day (rows).

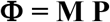

To construct the M and P matrices, we used a census interval of 5 days to calculate the proportion of flowers open for each individual within each 5-day window throughout the growing season. Since *L. salicaria* is hermaphroditic and self-incompatible, the same individuals are able to act as both pollen donors and pollen recipients

From the **Φ** matrix, we calculated the flowering synchrony index (*s*) following Weis et al. (2014) as the proportion of (co)variance explained by the dominant eigenvector. Specifically, *s* is bounded between 0 and 1, calculated as the first eigenvalue (*λ*_1_) of the **Φ** matrix divided by the sum of all eigenvalues 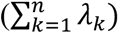 for **Φ**.

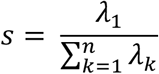

If all individuals have synchronized flowering, then there is only one mating pool, all mating pairs have the same mating probability, and *s* = 1. In self-incompatible species like *Lythrum salicaria*, the diagonal is a vector of zeros and *s* = 1 when all off-diagonal elements are equal. Alternatively, substructure in **Φ** representing separate mating pools will decrease *s* (Weis et al., 2014).

We also estimated the potential for assortative mating as the covariance between mates (*ρ*) for four phenotypic traits: (1) day of first flower, (2) size at flowering, (3) flowering duration, and (4) PC1. *ρ* for each trait was calculated following Weis et al. (2014):

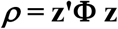

Where **z** is a vector of normalized phenotypic values (i.e., zero mean and unit variance) for a specific trait, and **z’** is its transpose. Here, it is explicit that nonzero values of *ρ* require phenotypic substructure of the matrix **Φ**. Therefore, a positive *ρ* for a given trait indicates asynchrony and assortment potential, with individuals tending to mate with others of similar phenotype. A negative *ρ* represents disassortment as individuals tend to mate with different phenotypes. Since both **z** and **Φ** contribute to *ρ*, values of *ρ* near 0 could represent synchronized mating, or asynchronized mating that is independent with respect to the phenotypic trait of interest.

We calculated **Φ,** *s* and *ρ* separately for each of the 20 populations in the experiment, as well as for all individuals in the common garden experiment. The latter simulates a large, genetically mixed population that might occur early in invasion, before locally adapted populations evolve. For each parameter in each population, we estimated confidence intervals by bootstrapping 1,000 random samples, with replacement, following Weis et al. (2014).

Because nonzero values of *ρ* require substructure in the **Φ** matrix, high flowering synchrony is expected to constrain *ρ* toward zero by homogenizing mating opportunities across phenotypes. Consistent with this expectation, Weis et al. (2014) demonstrated that populations and species with high flowering synchrony exhibit lower phenotypic correlation and therefore lower potential for assortment (Weis et al., 2014). To evaluate whether variation in flowering synchrony constrains assortative mating potential in our system, we fitted multiple linear regression models with *ρ* for each trait as a response variable and both *s* and latitude as continuous predictors. Positive values of *ρ* that do not covary with *s* indicate that assortative mating is present but is independent of the temporal overlap in flowering. While a significant association between *ρ* and latitude indicate that assortative mating is structured along a geographic gradient.

Assortative mating potential depends not only on the degree of asynchrony in flowering, but also on how phenotypes are mapped to mating probabilities in the **Φ** matrix. Since fitness and flowering time variation can shape these mating opportunities, and thereby influence flowering synchrony and the potential for assortment, we constructed four mating scenarios using our common garden data (Figure 2):

1. *Original field observation* (Figure 2A) refers to the raw data collected from the field, with individuals varying in flowering onset, duration, and total flower production over the growing season (Figure 2A).
2. *Equal fitness* (Figure 2B) refers to a fitness-standardized dataset, with all individuals producing the same total number of flowers without affecting variation in flowering schedule shape (Figure 2B). Here, we divided each individual’s flower counts by its total flower count across the growing season and then multiplied it by a factor of 100, simulating an equal flower output of 100 flowers across the season.
3. *Synchronized flowering onset* (Figure 2C) shifts flowering schedules so that all individuals start flowering at the same time but differ in fitness and flowering schedule shape (Figure 2C). Here, we used the un binned 5-day flower counts data and assumed that the first observation for every individual occurred in day one.
4. *Equal fitness with synchronized onset* (Figure 2D) combines the adjustments in 2 and 3 so that all individuals produce the same total number of flowers and start flowering on the same day but differ in the flowering schedule shape (Figure 2D).

**Figure 2.**
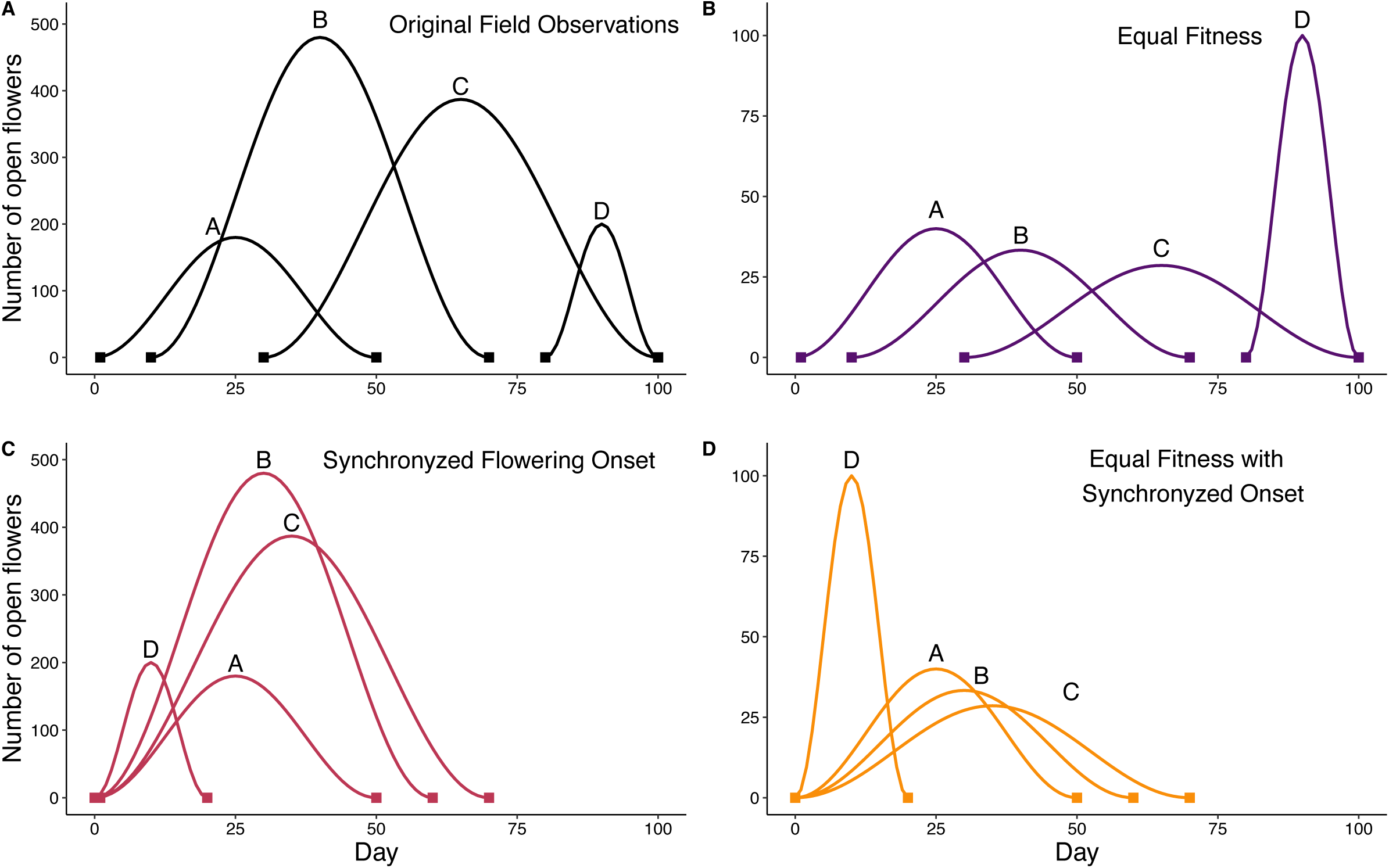
Conceptual model demonstrating variation in flowering schedules of four individuals (curves A, B, C, D). Pairwise mating probabilities are a function of (i) the area of overlap between two curves relative to (ii) the area of overlap with all other curves. To investigate contributions to assortative mating, flowering schedules can be compared across four scenarios: A) *Original field observations* use the schedules observed under natural growing conditions, with overlap determined by variation in flowering onset, curve shape, and total number of flowers produced. B) The *equal fitness* scenario controls for among-individual differences in total flower production. C) The *synchronized flowering onset* scenario controls for among-individual variation in the onset of flowering. D) *Equal fitness with synchronized onset* removes variation in both flowering onset and total flower number.

The un binned data used to create the *Synchronized flowering onset* and the *Equal fitness with synchronized onset* datasets had 38 missing flower counts from 35 individuals out of the 6,384 observations from 455 individuals. Therefore, we imputed these 38 missing values for the 35 individuals using a proximity matrix of the flowering trajectories across dates with the RandomForest package (Liaw & Wiener, 2002), with 10 iterations and 1000 trees. All data wrangling and statistical analyses scripts were performed in R studio (Posit team, 2025) and are available at (https://github.com/gomezquijano/AssortativeMatingL.salicaria).

## 3. Results

### 3.1 Flowering synchrony and the potential for assortment in the mixed population

To simulate an early invasion with mixed phenotypes, we analyzed the entire common garden experiment as a mixed population. Flowering synchrony in the *original field data* (see 1. *original field* scenario in Methods; Figure 2A) was intermediate (*s =* 0.50) and reduced relative to individual populations (Figure 3A). The potential for assortative mating based on phenotypic correlations between mates ranged from 0.15 to 0.28 (Figures 3B-E). Day of first flower (*ρ* = 0.28; Figure 3B) and PC1 (*ρ* = 0.26; Figure 3E) exhibited the highest potential for correlated mating, with less assortment potential for size at flowering (*ρ* = 0.15; Figure 3C) and flowering duration (*ρ* = 0.15; Figure 3D).

**Figure 3.**
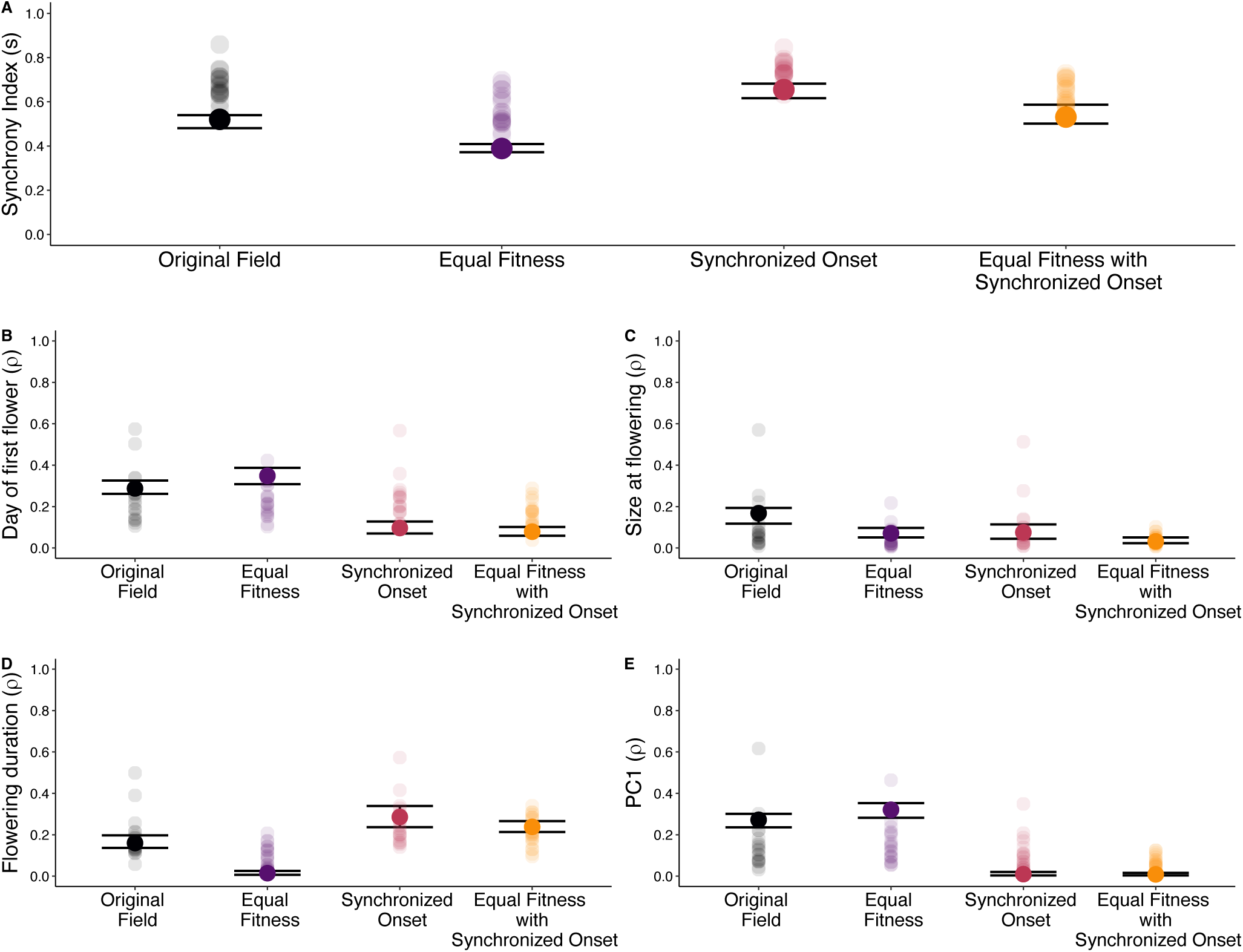
Flowering synchrony index (A) and potential for assortative mating (*ρ*) by day of first flower (B), size at flowering (C), flowering duration (D) and PC1 as a composite measure of day of first flower and size at flower (E), each calculated for individuals grouped into 20 populations (translucent circles) or the entire mixed population of the common garden (N = 455 individuals; solid circles). Values are calculated for each of four different scenarios: the *original field observations* (black), adjusted for *equal fitness* (purple), adjusted for *synchronized flowering onset* (red) and adjusted for *both equal fitness and flowering onset* (orange). Horizontal bars show 95% confidence intervals (CI) for the mixed population, based on 1,000 random samples, with replacement.

When individuals within the mixed population were standardized to the same number of flowers (see 2. *Equal fitness* scenario in Methods; Figure 2B), flowering synchrony decreased significantly (*s =* 0.39; Figure 3A) relative to the *original field* observations, as did the potential for assortative mating by size at flowering (*ρ =* 0.07; Figure 3C) and flowering duration (*ρ =* 0.014; Figure 3D). In contrast, adjusting for equal fitness increased potential for assortment by day of first flower (*ρ =* 0.35; Figure 3C) and PC1 (*ρ =* 0.32; Figure 3C).

When flowering time was equalized across individuals (see 3. *Synchronized flowering onset* scenario in Methods; Figure 2C), flowering synchrony (*s =* 0.65) increased significantly relative to the three other scenarios (Figure 3A), as did assortative mating potential by flowering duration (*ρ =* 0.28; Figure 3D). In contrast, the phenotypic correlation between mates for day of first flower (*ρ =* 0.09; Figure 3B), size at flowering (*ρ =* 0.07; Figure 3C), and PC1 (*ρ =* 0.01; Figure 3C) all declined relative to the *original field* dataset.

Finally, when both fitness and flowering onset were adjusted (see 4. *Equal fitness with synchronized onset* in Methods; Figure 2D), synchrony in the mixed population was moderate (*s =* 0.52) and similar in magnitude to the *original field* data (Figure 3A). The potential for assortative mating by flowering duration was also elevated (*ρ =* 0.24; Figure 3D), whereas day of first flower (*ρ =* 0.08; Figure 3B), size at flowering (*ρ =* 0.03; Figure 3C) and PC1 (*ρ =* 0.008; Figure 3C) were the lowest among all scenarios.

### 3.2 Flowering synchrony and the potential for assortment in individual populations

In the analysis of the *original field data*, flowering synchrony of the 20 individual populations ranged from 0.56 to 0.86, each of these values was higher than the synchrony value of *s =* 0.50 calculated for the entire mixed population (Figure 3A). With a few exceptions, the potential for assortative mating was generally lower in individual populations compared to the entire mixed population, whether calculated by day of first flower (*ρ* = 0.11 to 0.57; Figure 3B), size at flowering (*ρ* = 0.09 to 0.57; Figure 3C), flowering duration (*ρ* = 0.06 to 0.50; Figure 3D), or PC1 (*ρ* = 0.03 to 0.62; Figure 3E).

For the *equal fitness* scenario, flowering synchrony among populations ranged from 0.39 to 0.69, and was generally higher than the *s* = 0.39 calculated for the entire mixed population (Figure 3A). Similar to the *original field* observations, the potential for assortative mating calculated for each individual population was typically lower than in the mixed populations for day of first flower (*ρ* = 0.1 to 0.42, Figure 3B), size at flower (*ρ* = 0.05 to 0.21, Figure 3C) and PC1 (*ρ* = 0.05 to 0.46, Figure 3E). Conversely, the phenotypic correlations among mates for flowering duration were higher for individual populations (*ρ* = 0.03 to 0.21), when compared to the overall mixed population (*ρ* = 0.15) (Figure 3D).

In the *synchronized flowering onset* scenario, synchrony of individual populations ranged from 0.63 to 0.85, with all but one population being higher than the metapopulation *s* = 0.65 (Figure 3A). Potential for assortative mating was more variable but generally higher by day of first flower (*ρ* = 0.09 to 0.57; Figure 3B) and PC1 (*ρ* = 0.03 to 0.35; Figure 3E), and lower for size at flowering (*ρ* = 0.009 to 0.51, Figure 3C) and flowering duration (*ρ* = 0.14 to 0.57, Figure 3D).

Finally, for the *equal fitness with standardized onset* scenario, synchrony was similar in magnitude to the *original field observations* but less variable among populations (*s* = 0.53 to 0.73; Figure 3A). Compared to the mixed population, individual populations typically had a higher potential for assortative by day of first flower (*ρ* = 0.04 to 0.29, Figure 3B) and PC1 (*ρ* = 0.01 to 0.13, Figure 3E). In contrast, variability among populations in potential assortment by size at flowering (*ρ* = 0.007 to 0.1, Figure 3C) and flowering duration (*ρ* = 0.09 to 0.34, Figure 3D) overlapped with the mixed population at intermediate values.

### 3.3 Clines in flowering synchrony and the potential for assortment

Among-population variation in synchrony and assortative mating potential by size at flowering covaried with latitude (Figure 4) but no other traits did (Supplemental Figure 1). In the *original field observations*, synchrony index decreased with latitude (*β* = − 0.01), with higher synchrony in southern populations (Table 1A; Figure 4A). There was also a quadratic relationship between latitude and potential for assortative mating by size at flowering (*β* = 0.260) due to elevated assortment at the most southern latitudes (Table1A; Figure 4A). However, in the *equal fitness* scenario, flowering synchrony had a quadratic relationship (*β* = − 0.17), with increased synchrony at intermediate latitudes (Table 1B; Figure 4B), but no clinal variation in the potential for assortative mating. In the *synchronized flowering onset* scenario, synchrony exhibited a quadratic association with latitude (*β* = −0.01) similar to the *equal fitness* scenario, but the assortative mating by size at flowering (*β* = 0.26), was more similar to the *original field observations* (Table 1C; Figure 4C). Lastly, we found no evidence of clinal variation in synchrony or in the potential for assortative mating in the *equal fitness with standardized onset* scenario (Table 1D; Figure 4D).

**Figure 4.**
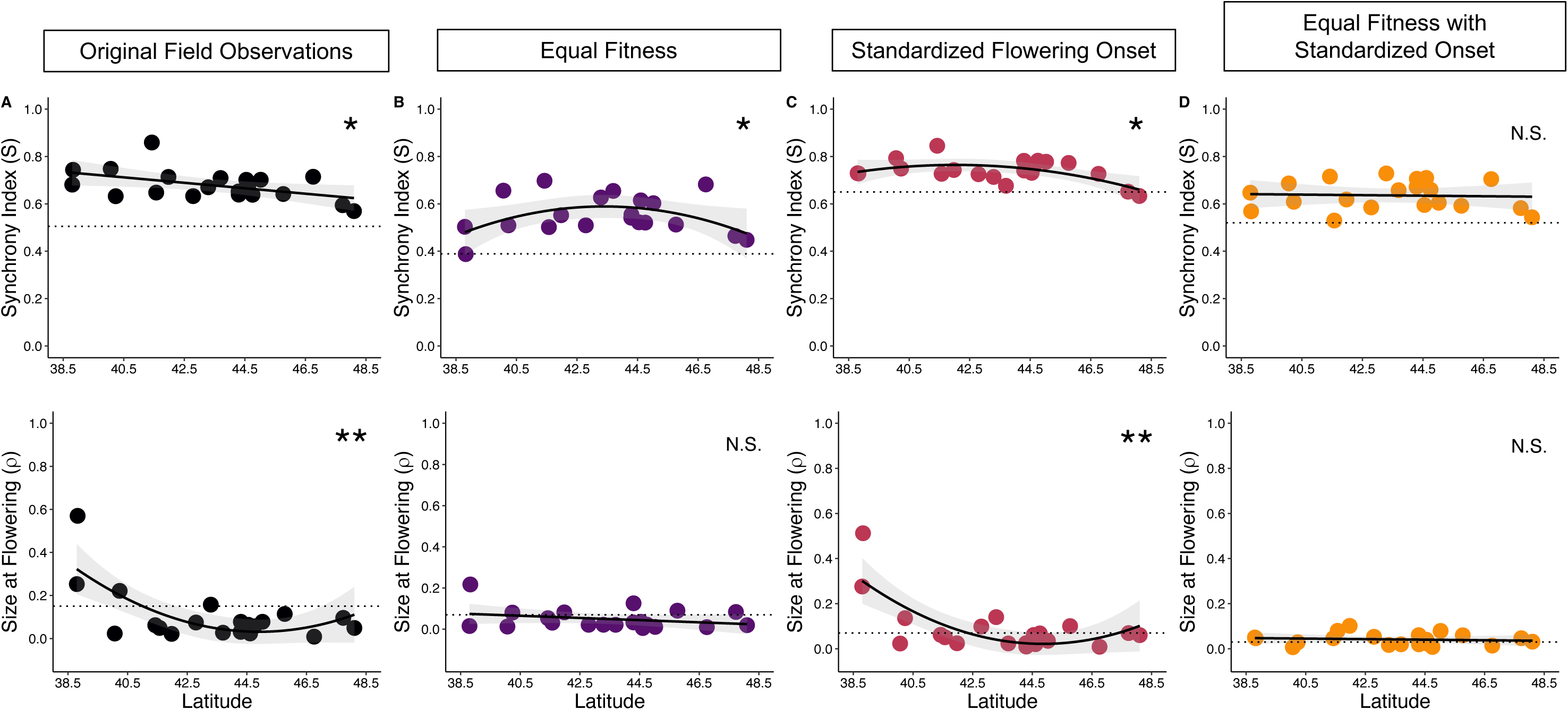
Bivariate plots showing variation in flower synchrony (*s*, top row) and the potential for assortative mating for size at flowering (*ρ*, bottom row) calculated for 20 locally adapted populations (dots) of *L. salicaria* sampled along a latitudinal gradient and grown in a common garden at mid-latitude. Columns show calculations for each scenario outlined in Figure 2: A) *Original field data* (black). B) *Equal fitness* (purple). C) *Synchronized flowering onset* (red). D) *Equal fitness with synchronized onset* (orange). Regression lines are fitted using linear or quadratic regression models, with best model determined AIC scores and with significance denoted as * p < 0.05, ** p < 0.01 or *** p < 0.001. Horizontal dotted lines show the calculation for the entire mixed population of the common garden, equal to the mixed population means shown in Figure 3.

**Table 1.**
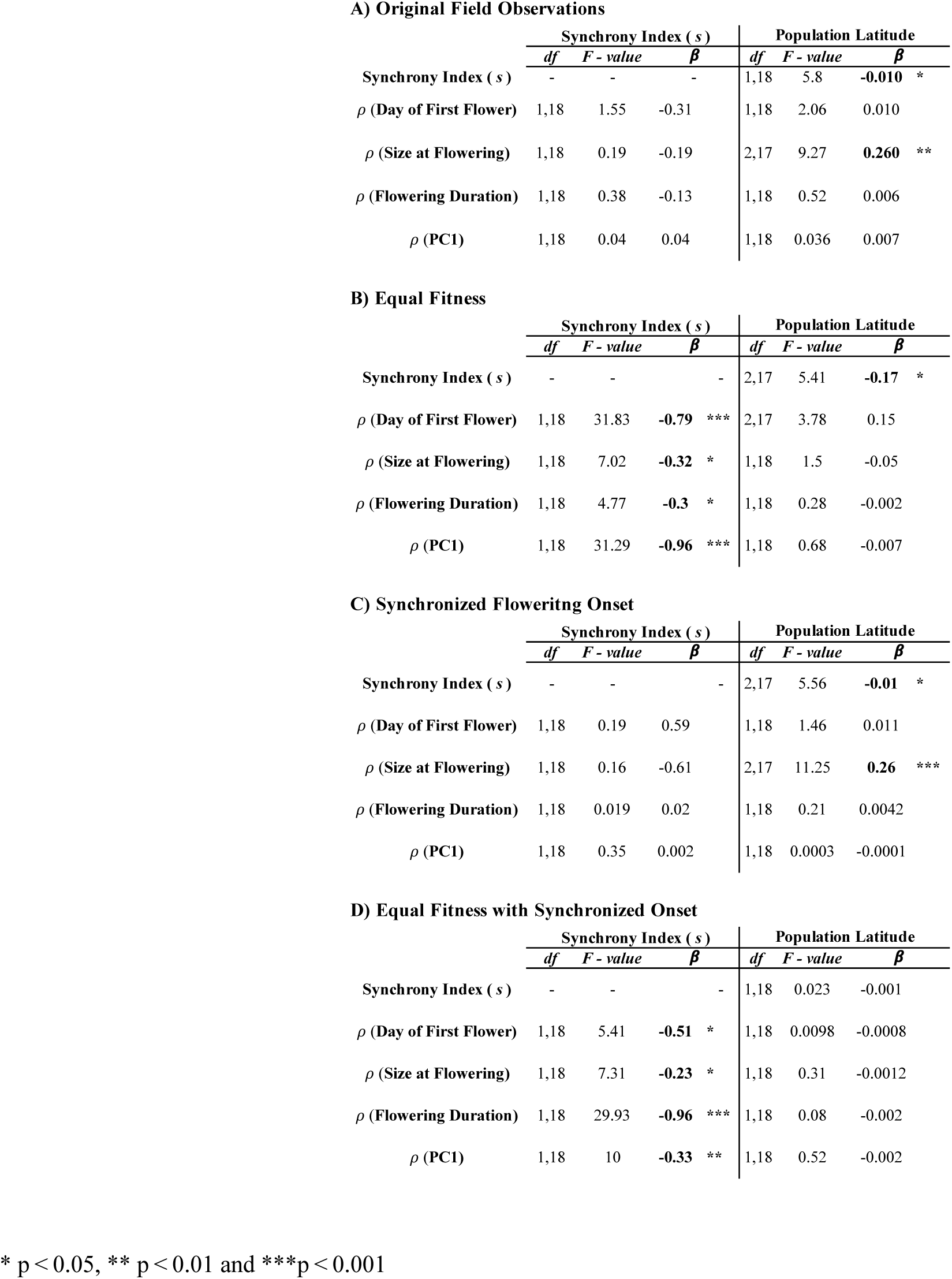
Regression statistics predicting the potential for assortative mating for flower time, size at flowering, flowering duration and PC1 from population-level flower synchrony indexes and latitude. Regression models were performed on A) *Original field observations* B) *equal fitness* C) *synchronized flowering onset* and D) *Equal fitness with synchronized onset* datasets. Dashes represent predictors that were not included in the model. For a detailed description of each dataset see section 2.4. Bold values represent significance of the predictor variable (columns) at p < 0.05.

| **A) Original Field Observations** |  |  |  |  |  |  |
| --- | --- | --- | --- | --- | --- | --- |
|  | Synchrony Index ( *s* ) |  |  | Population Latitude |  |  |
| | *df* | *F - value* | $\beta$ | *df* | *F - value* | $\beta$ |
| Synchrony Index ( *s* ) | - | - | - | 1,18 | 5.8 | **-0.010** * |
| $\rho$ (Day of First Flower) | 1,18 | 1.55 | -0.31 | 1,18 | 2.06 | 0.010 |
| $\rho$ (Size at Flowering) | 1,18 | 0.19 | -0.19 | 2,17 | 9.27 | **0.260** ** |
| $\rho$ (Flowering Duration) | 1,18 | 0.38 | -0.13 | 1,18 | 0.52 | 0.006 |
| $\rho$ (PC1) | 1,18 | 0.04 | 0.04 | 1,18 | 0.036 | 0.007 |
| **B) Equal Fitness** |  |  |  |  |  |  |
|  | Synchrony Index ( *s* ) |  |  | Population Latitude |  |  |
| | *df* | *F - value* | $\beta$ | *df* | *F - value* | $\beta$ |
| Synchrony Index ( *s* ) | - | - | - | 2,17 | 5.41 | **-0.17** * |
| $\rho$ (Day of First Flower) | 1,18 | 31.83 | **-0.79** *** | 2,17 | 3.78 | 0.15 |
| $\rho$ (Size at Flowering) | 1,18 | 7.02 | **-0.32** * | 1,18 | 1.5 | -0.05 |
| $\rho$ (Flowering Duration) | 1,18 | 4.77 | **-0.3** * | 1,18 | 0.28 | -0.002 |
| $\rho$ (PC1) | 1,18 | 31.29 | **-0.96** *** | 1,18 | 0.68 | -0.007 |
| **C) Synchronized Flowering Onset** |  |  |  |  |  |  |
|  | Synchrony Index ( *s* ) |  |  | Population Latitude |  |  |
| | *df* | *F - value* | $\beta$ | *df* | *F - value* | $\beta$ |
| Synchrony Index ( *s* ) | - | - | - | 2,17 | 5.56 | **-0.01** * |
| $\rho$ (Day of First Flower) | 1,18 | 0.19 | 0.59 | 1,18 | 1.46 | 0.011 |
| $\rho$ (Size at Flowering) | 1,18 | 0.16 | -0.61 | 2,17 | 11.25 | **0.26** *** |
| $\rho$ (Flowering Duration) | 1,18 | 0.019 | 0.02 | 1,18 | 0.21 | 0.0042 |
| $\rho$ (PC1) | 1,18 | 0.35 | 0.002 | 1,18 | 0.0003 | -0.0001 |
| **D) Equal Fitness with Synchronized Onset** |  |  |  |  |  |  |
|  | Synchrony Index ( *s* ) |  |  | Population Latitude |  |  |
| | *df* | *F - value* | $\beta$ | *df* | *F - value* | $\beta$ |
| Synchrony Index ( *s* ) | - | - | - | 1,18 | 0.023 | -0.001 |
| $\rho$ (Day of First Flower) | 1,18 | 5.41 | **-0.51** * | 1,18 | 0.0098 | -0.0008 |
| $\rho$ (Size at Flowering) | 1,18 | 7.31 | **-0.23** * | 1,18 | 0.31 | -0.0012 |
| $\rho$ (Flowering Duration) | 1,18 | 29.93 | **-0.96** *** | 1,18 | 0.08 | -0.002 |
| $\rho$ (PC1) | 1,18 | 10 | **-0.33** ** | 1,18 | 0.52 | -0.002 |

\* $p < 0.05$ , \*\* $p < 0.01$ and \*\*\* $p < 0.001$

### 3.4 Covariances of flowering synchrony and the potential for assortment

Flower synchrony predicted phenotypic correlations between mates for all traits only in scenarios where individual fitness was equalized. In the *original field observations* (*β* = − 0.31 to 0.04; Table 1C; Figure 5C) and *synchronized flowering onset* (*β* = − 0.96 to − 0.3; Table 1C; Figure 5C) scenarios synchrony indexes (*s*) were not significant predictors of phenotypic correlations between mates (*ρ*) for any of the measured traits. Meanwhile, for the *equal fitness* scenario, synchrony indexes were significantly and negatively associated with phenotypic correlations between mates (*ρ*) for all traits (*β* = − 0.61 to 0.59; Table 1B; Figure 5B). When both flower number and flowering time were equalized, in the *equal fitness with synchronized onset* scenario, synchrony indexes were significantly and negatively associated with phenotypic correlations between mates for all traits (*β* = − 0.23 to 0.96), indicating that greater asynchrony (low *s* values) increase the potential for assortative mating (Table 1D; Figure 5D).

**Figure 5.**
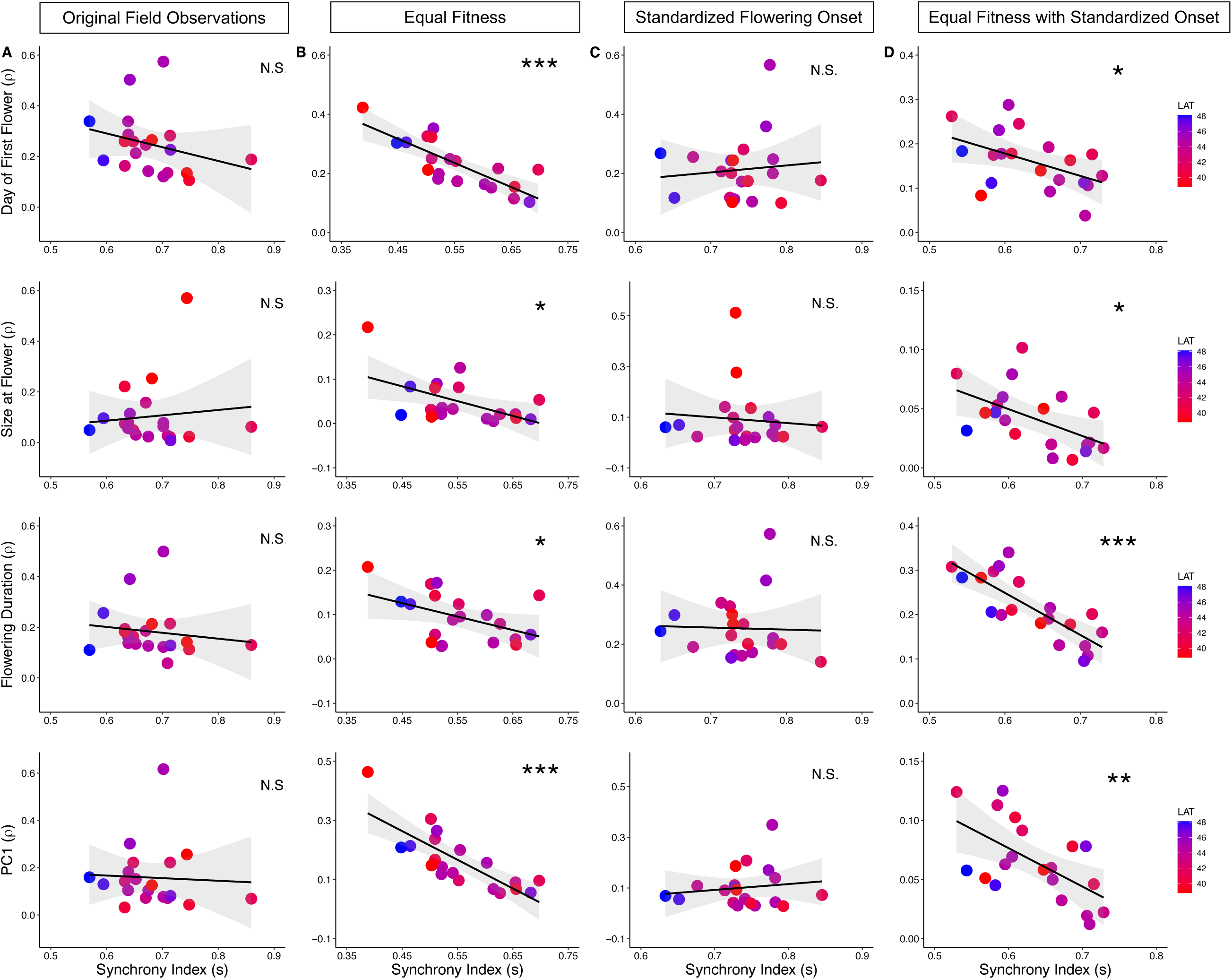
Predictive relationship of synchrony indexes (*s*) and latitude on the potential of phenotypic assortative mating (*ρ*). Phenotypic correlations between mates (*ρ*) for (top to bottom) day of first flower, size at flowering, flowering duration and PC1 plotted against population synchrony indexes (*s*) under A) *Original field data* conditions, B) *equal fitness*: equal number of flowers produced over the season, C) *synchronized flowering onset*: equal flowering onset for all individuals and, D) *equal fitness with synchronized onset:* equal flowering onset and number of flowers produced over the season for all individuals. Each circle represents a population, regression lines are fitted using linear regression models and populations are color coded based on latitudinal origin, with warmer tones representing lower latitudes and cooler tones representing higher latitudes. asterisks represent significance of synchrony index as a predictor variable at * p < 0.05, ** p < 0.01 and ***p < 0.001. Significance of latitude as a predictor is shown in Table 1, Figure 4 and Supplementary Figure 1.

## 4. Discussion

Reduced overlap in flowering schedules creates asynchrony in mating probabilities that partition populations into smaller mating pools, promoting phenological assortment that can accelerate the rate of stochastic and adaptive (Crow & Felsenstein, 1968; Weis & Kossler, 2004; Wright, 1921). Yet, the contribution of phenological assortative mating to contemporary evolution remains largely unexplored. Using a common garden experiment, we explored the importance of assortment during invasion by creating an artificially mixed population from seed families sampled across 20 *Lythrum salicaria* populations spanning 10 degrees of latitude in eastern North America (Figure 1). In contrast to any individual natural population, our artificially mixed population is more similar to an early invasion scenario characterized multiple introductions from genetically diverse origins, before populations diverge due to natural selection and drift. To explore the potential for assortment to accelerate adaptive evolution during invasion, we quantified flowering schedule synchrony (*s*) and the resulting phenotypic correlations between mates (*ρ*) (Weis et al., 2014) for the mixed and individual populations. We found that mixed population were less synchronous relative to locally adapted populations, which also exhibited latitudinal clines in *s* and *ρ* for size at flowering. Furthermore, the predicted negative correlation between *s* and *ρ* was masked by among-individual fitness variation in the common garden. Below we discuss the factors affecting assortative mating potential in *L*. *salicaria* and more generally how these same factors can affect the rate of adaptive evolution during biological invasions.

### 4.1 Insights from a mixed population: Flowering schedule variation, potential for assortment and effects of fitness and flowering time in an early invasion scenario

Flowering synchrony emerges from joint variation in flowering onset, duration, and flower production which structures mating opportunities and the potential for phenological assortative mating. Because flower production strongly affects both siring success (Austen & Weis, 2016) and temporal structure (Weis & Kossler, 2004), removing fitness variation by equalizing flower production (*equal fitness*) reduced flowering synchrony (*s =* 0.39) relative to the *original field observations* (*s =* 0.50; Figure 3A); revealing the presence of mating dominance from high fitness individuals. In contrast, as expected removing flower onset variation (*synchronized flowering onset*) increased flower schedule synchrony (*s =* 0.65) (Figure 3A). Together these indicate that while differences in flowering time are a major driver of mating structure (Fox, 2003; Weis & Kossler, 2004), variation in fitness components can also substantially reshape mating opportunities. When matings are dominated by a few highly fecund individuals, increased flowering schedule overlap can elevate synchrony and weaken assortative mating potential. This can cause an increase in effective gene flow among phenotypically dissimilar individuals, potentially reducing phenotypic variance and constraining adaptive responses to selection.

Although both flower number (Lyons & Mully, 1992) and flowering onset (Ison et al., 2014; Weis et al., 2014) are known to exhibit assortative mating potential in natural populations, their relative strength depends on how mating opportunities are structured. Under the *original field observations* assortative mating potential was highest for day of first flower (*ρ* = 0.28; Figure 3B) and PC1 (*ρ* = 0.26; Figure 3E). Removing variation in flower production (*equal fitness*) increased assortment potential for first flower (*ρ =* 0.35; Figure 3B) and PC1 (*ρ =* 0.32; Figure 3E), while equalizing flowering onset (*synchronized flowering onset*) drastically reduced assortative mating potential for these two traits to *ρ =* 0.07 and *ρ =* 0.01 respectively (Figure 3B and 3E). Such differences highlight day of first flower as the major axis of temporal assortment (Fox, 2003; Weis et al., 2014) and suggests that mating dominance by high-fitness individuals could decrease assortment potential in mixed populations during early invasion stages. Yet, the persistence of positive assortment potential through traits such as flowering duration (Figure 3D) in the absence of flowering onset variation, indicates that non-random mating can remain flexible when temporal variation is constrained. This context dependent and flexible pattern aligns with the strong genetic correlations among life-history traits and substantial standing genetic variation for individual traits in *L. salicaria* (Colautti & Barrett, 2011), suggesting that assortative mating can emerge across multiple correlated traits (Weis, 2005) in ways that help preserve phenotypic variance for rapid adaptive responses to occur.

Assortative mating in plants has been shown to arise through covariance in traits associated with mating opportunities such as stem height and stem diameter, rather than flowering phenology alone (Weis, 2005). In *L. salicaria* positive mate covariances (*ρ*) for flowering duration (Figure 3C) and size at flower (Figure 3D), while weaker than for day of first flower (Figure 3B) and PC1 (Figure 3E), are particularly relevant given the strong trade-offs between flowering onset and size (Colautti & Barrett, 2013) despite weak genome-wide neutral differentiation among locally adapted populations (Gómez Quijano, 2026). Because assortative mating acts on phenotypic similarity rather than on genetic identity (Wright, 1921), recombination can maintain high admixture at neutral loci while increasing homozygosity at phenology associated traits (Crow & Felsenstein, 1968). In this way, assortative mating may promote adaptive divergence in key life-history traits without requiring broad neutral structure (Crow & Felsenstein, 1968), which could explain why locally adapted population of *L. salicaria* maintain high levels of genome wide similarities.

### 4.2 Insights from locally adapted populations: Clinal flowering schedule variation, potential for assortment and effects of fitness and flowering time in contemporary populations

Assortative mating can persist within locally adapted populations even when flowering schedules overlap. Because directional or stabilizing selection is expected to reduce phenotypic variance for traits under selection (Lande & Arnold, 1983), locally adapted populations should exhibit greater flower synchrony than mixed populations. In contrast, mixed populations, particularly during early invasion, are expected to retain elevated trait variance because selection and drift have had limited time to purge maladaptive phenotypes (Dlugosch & Parker, 2008). Consistent with these expectations, flower synchrony was higher in individual populations than in the mixed population and reached values as high as 0.86 (Figure 3A). Although phenotypic correlations between mates for day of first flower (Figure 3B), size at flowering (Figure 3C), and PC1 (Figure 3E) were often lower in individual populations than in the mixed population, assortative mating potential remained positive. Phenotypic correlations between mates across all traits and scenarios were greater than zero reaching values as high as 0.62 (Figure 3B-E), indicating that locally adapted populations retain sufficient phenotypic variation in traits influencing mating opportunities (Ison et al., 2014). The persistent of positive assortment potential despite extensive flowering overlap within populations, also suggests the presence of “cryptic” assortative mating, in which mating bias emerges from day-to-day variation in flower frequency rather than from temporal separation (Fox, 2003). Further supporting this phenomenon is the *standardized flowering onset* scenario, for which populations despite being highly synchronous (*s* = 0.63 to 0.85; Figure 3A) maintained positive mate covariances (*ρ* = 0.03 to 0.57). Therefore, “cryptic” assortative mating (Fox, 2003) driven by fitness variation, may help maintain adaptive potential despite local adaptation, facilitating future evolutionary responses in natural populations.

The persistence of variation in flower synchrony and assortative mating potential within locally adapted populations, suggests these processes may contribute to the evolution of geographic clines in adaptive trait differentiation. Environmental variation exerts strong selection pressures on adaptive traits leading to the formation of clines in phenology, size, and fitness (Barton, 1999; Colautti & Barrett, 2013; Soularue & Kremer, 2012; Yang et al., 2021). Such differentiation is often reinforced by reduced maladaptive gene flow among populations (Fitzpatrick & Reid, 2019) facilitated by phenological schedule asynchrony and assortative mating (Soularue & Kremer, 2012). Consistent with this expectation, both the *original field* and the *synchronized flowering onset* scenarios showed that higher latitude populations were more asynchronous (Table 1; Figure 4A) but exhibited weaker mate covariance for size at flowering (Figure 4B). This likely reflects earlier flowering, longer flowering durations (Akbar et al., 2025) and, reduced phenotypic variance for size at flowering at higher latitudes. However, in *L. salicaria,* geographic structure in assortment potential may be mediated by fitness-dependent differences in mating opportunities. Latitudinal clines in *ρ* for size at flowering disappeared when flower number, a proxy for fitness, was standardized (*equal fitness* and *equal fitness with synchronized onset*; Table 1; Figure 4). Preventing high-fitness individuals within each population from disproportionately dominating matings, eliminated clinal variation in assortment potential. Highlighting within population fitness variation as a key contributor of the evolution and maintenance of trait-specific geographic clines in assortative mating potential.

### 4.3 Effects of fitness and flowering time variation on the expected relationship between flowering schedule synchrony and phenotypic correlations between mates

Phenological assortative mating potential does not appear to be solely determined by flowering onset, instead it is strongly mediated by fitness-weighted mating opportunities. Assortative mating in plants is dependent on the temporal structuring of mating opportunities (Fox, 2003). Therefore, when individuals differ in their flowering schedules, reproductive asynchrony and assortment potential are expected to increase (Weis & Kossler, 2004; Weis at al. 2014). However, unlike Weis at al. (2014), we found no significant association between *s* and *ρ* in the *original field* or the *synchronized flowering onset* scenarios (Table 1; Figure 5A,C). The absence of this relationship is to be expected in the *synchronized flowering onset* scenario, where populations are expected to have greater flowering schedule synchrony and reduced temporal structuring. Nonetheless, the nonsignificant relationship between reproductive synchrony and assortment potential was unexpected in the *original field observations*, where individuals experience variation in flowering onset and flower output. This phenomenon could be explained by two hypotheses: first, that low flowering schedule synchrony (mating asynchrony) is necessary but not a sufficient cause for phenological assortative mating for a given trait (Weis at al. 2014). And second, that within locally adapted populations, “cryptic” assortative mating may be driven by variation in flower output (Fox, 2003). By allowing a small number of highly fecund individuals to contribute disproportionately to matings, fitness variation can maintain both high flowering synchrony and positive mate covariances, thereby masking the expected synchrony and assortment potential relationship. This is further supported by the *equal fitness* and *equal fitness and standardized onset* scenarios where the significant negative relationship between *s* and *ρ* emerges only after fitness differences are removed (Table 1; Figure 5B,D).

Together, these results suggest that within smaller locally adapted populations of *L. salicaria*, fitness variation can structure mating opportunities in ways that allow assortative mating potential to persist despite high flowering schedule synchrony. During invasion, if the high levels of standing genetic variation generated by multiple introductions from the native range contributes to differences in reproductive output among individuals (Chun et al., 2009; Dlugosch & Parker, 2008; Keller & Taylor, 2010), fitness-weighted mating opportunities could maintain positive mate covariances and reduce effective gene flow among individuals. As a consequence, additive genetic variance for phenology associated traits is likely to be preserved within the population, facilitating rapid adaptive evolution even under high admixture scenarios.

### 4.4 Caveats

Our study serves as a proxy to investigate how phenological synchrony could promote assortative mating in natural populations and how this mechanism may have also facilitated local adaptation during early invasion. In our study, flower synchrony in the mixed common garden population (*original field observations*; Figure 3A), was lower than reported for most tree and herbaceous species (Augspurger, 1983; Elzinga et al., 2007; Mahoro, 2002; Weis et al., 2014). Meanwhile, *ρ* values were comparable to those reported for day of first flower in natural populations of other Ontario field species (Weis et al., 2014). Despite these similarities, several caveats should be considered when interpreting our results. First, the common garden represents a reconstruction of the phenotypic variation that may have existed during the early stages of invasion rather than a direct observation of historical populations. Consequently, our estimates of flowering synchrony and assortative mating potential should be viewed as approximations of the mating opportunity structure that could have emerged following admixture from multiple introduction events. Second, although flowering overall is a prerequisite for mating, in a self-incompatible system like *L. salicaria*, pollen transfer depends on pollen production and pollinator behavior and abundance (Harder & Barrett, 1996), factors that were not incorporated into our framework.

Finally, our approach assumes that all individuals have equal access to potential mates, however *L. salicaria* exhibits a tristylous mating system in which mating is constrained by floral morph compatibility (Darwin, 1877; O’Neil, 1994). Incorporating morph frequencies into estimates of mating opportunities would provide a more realistic representation of realized mating patterns. While doing so would require substantial modification of the mating matrix framework and would considerably reduce the sample size available for the analysis, it represents an important avenue for future work. Likewise, incorporating measures of seed production, pollen production, pollinator abundance and behaviour could help bridge the gap between potential and realized matings as well as potential and effective gene flow. Nevertheless, because flowering schedule overlap defines the temporal window in which compatible matings can occur, our results provide valuable insight into how phenological variation and fitness differences may structure mating opportunities and influence the potential for assortative mating of natural populations during early invasion and across latitudinal gradients.

## 5. Conclusion

Although population differentiation and adaptation are driven by selection, drift and gene flow (Dobzhansky, 1937; Mayr, 1963; Wright, 1931), assortative mating should be considered as a significant complementary process capable of shaping the outcomes of such forces and influence evolutionary trajectories. Using a mixed population in a common garden as an analogue for early invasion, we showed that individual flowering schedule variation can promote strong assortative mating potential by structuring mating opportunities through both temporal and fitness-mediated differences. Moreover, we demonstrate how even though, temporal overlap establishes the opportunity for mating (Weis & Kossler, 2004), the strength of assortment depend on how reproductive opportunities are distributed among individuals, supporting that while important, flowering synchrony alone is not sufficient to determine assortative mating potential (Weis et al., 2014). In *L. salicaria* fitness-weighted mating opportunities arising from variation in flower production can maintain positive mate covariances even when flowering schedules overlap extensively, revealing a form of “cryptic” assortative mating (Fox, 2003) that would have not be predicted from flowering schedule synchrony estimates alone.

Our study has important implications for understanding rapid adaptation from standing genetic variation. During invasion, admixture among multiple introductions can generate high standing genetic variation, increasing opportunities for rapid evolutionary responses while simultaneously increasing the probability of maladaptive allele introduction through gene flow (Dlugosch & Parker, 2008). By biasing mating toward phenotypically similar and highly fecund individuals, assortative mating may reduce effective gene flow among distinct phenotypes (Devaux & Lande, 2008), increasing phenotypic variance (Wright, 1921) and helping maintain adaptive trait combinations despite extensive admixture (Godineau et al., 2022). More broadly, our results suggest that non-random mating may contribute to the maintenance of phenotypic variation, the formation of geographic clines and the persistence of local adaptation across environmental gradients and distinct selection regimes. Ultimately, as climate change continues to alter reproductive phenology, the interaction between non-random mating with adaptive and stochastic evolutionary forces may play an increasingly important role in determining the evolutionary fates of natural populations.

## Data Availability

Reproducible code for all analyses and figures is available at: https://github.com/gomezquijano/AssortativeMatingL.salicaria, flower census data is available at: [insert dryad]

## Author Contributions

MJGQ and RIC designed the study; MJGQ led the data analysis and manuscript writing with support from RIC. The study was designed based on filed data collected by MNA, EC and MJGQ and preliminary analyses by MNA and EC.

## Funding

[insert]

## Conflict of Interests

The authors declare no conflict of interest

## Acknowledgements

The authors acknowledge Matthew Kleronomos for assistance with field data collection and the Queen’s University Biological station (QUBS) especially senior manager, Sonia Nobrega, for providing the facilities and resources needed to establish and maintain the common garden.

## Notes

### Competing Interest Statement

The authors have declared no competing interest.

