## Supplementary Material for "Assortative mating may have facilitated rapid evolution of locally adaptive phenologies during a biological invasion"

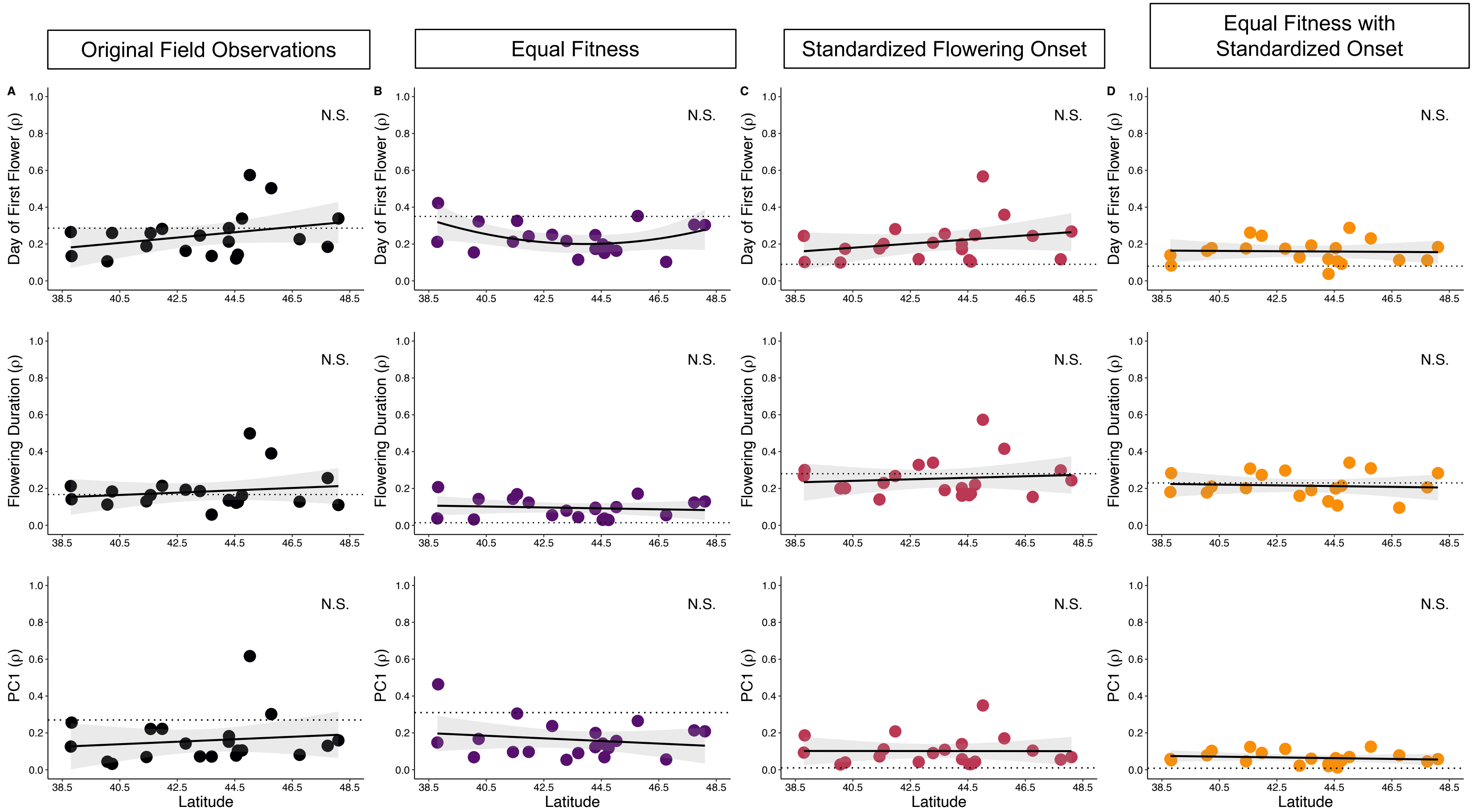


**Supplementary Figure 1.** Non-significant latitudinal clines in the potential for assortative mating for day of first flower (top), flowering duration (middle) and PC1 (bottom) of 20 locally adapted populations of *L. salicaria.* correlation between mates for each trait is calculated for populations under A) *Original field data* conditions (black), B) *equal fitness* (purple), C) *synchronized flowering onset* (red) and D) *equal fitness with synchronized onset* (orange). Each population across the gradient is represented by a circle, regression lines are fitted using linear or polynomial (quadratic) regression models. Dotted line represents the metapopulation average shown in Figure 3.
